# Decoupling the impact of microRNAs on translational repression versus RNA degradation in embryonic stem cells

**DOI:** 10.1101/316679

**Authors:** Jacob W. Freimer, TJ Hu, Robert Blelloch

## Abstract

Translation and mRNA degradation are intimately connected, yet the mechanisms that regulate them are not fully understood. Here we studied the link between translation and mRNA stability in embryonic stem cells (ESCs). Transcripts showed a wide range of stabilities, which correlated with their translation levels. The protein DHH1 links translation to mRNA stability in yeast; however loss of the mammalian homolog, DDX6, in ESCs did not disrupt the correlation across transcripts. Instead, the loss of DDX6 led to upregulated translation of microRNA targets, without concurrent changes in mRNA stability. The *Ddx6* knockout cells were phenotypically and molecularly similar to cells lacking all microRNAs (*Dgcr8* knockout ESCs). These data show that the loss of DDX6 can separate the two canonical functions of microRNAs: translational repression and transcript destabilization. Furthermore, these data uncover a central role for translational repression independent of transcript destabilization in defining the downstream consequences of microRNA loss.

## INTRODUCTION

Gene expression is determined through a combination of transcriptional and post-transcriptional regulation. While transcriptional regulation has been extensively studied, less is known about how the regulation of mRNA stability contributes to overall mRNA levels. Mammalian mRNAs display a wide range of half-lives ranging from minutes to over a day (Schwanhäusser et al., 2011). The wide range of mRNA stabilities are regulated by both intrinsic sequence features as well as the binding of regulatory factors such as microRNAs and RNA-binding proteins (Cheng, Maier, Avsec, Rus, & Gagneur, 2016; Hasan, Cotobal, Duncan, & Mata, 2014; Wu & Brewer, 2012). However, the identity of such features and regulatory factors as well as how they impact mRNA stability are not well understood.

One process that is intimately linked to mRNA decay is translational repression (Roy & Jacobson, 2013). Quality control mechanisms such as nonsense mediated decay, no go decay, and non stop decay sense aberrant translation and lead to mRNA degradation (Parker, 2012; Shoemaker & Green, 2012). Translational repression and mRNA decay are also linked independent of either a quality control or a stress response (Huch & Nissan, 2014). In yeast, inhibition of translation initiation through either 5’ cap binding mutants or drug treatment leads to accelerated mRNA decay (Chan, Mugler, Heinrich, Vallotton, & Weis, 2017; Huch & Nissan, 2014; Schwartz & Parker, 1999). Conversely treatment with cycloheximide, which blocks ribosome elongation, stabilizes mRNAs (Beelman & Parker, 1994;Chan et al., 2017; Huch & Nissan, 2014). How upstream factors influence mRNA stability through the regulation of translation initiation and elongation is poorly understood.

Whether microRNAs (miRNAs) primarily impact translation or mRNA degradation has been intensely debated (Iwakawa & Tomari, 2015; Jonas & Izaurralde, 2015). MiRNAs are small, non-coding RNAs that bind to the 3’ UTR of their target transcripts to inhibit translation and/or induce mRNA destabilization (Fabian & Sonenberg, 2012; Jonas & Izaurralde, 2015). The impact of miRNAs on the translation and expression of endogenous transcripts has been measured using ribosome profiling which measures the ratio of ribosome protected fragments to input mRNA (this ratio is termed translational efficiency) and RNA-Seq which measures RNA levels. Simultaneous ribosome profiling and RNA-Seq experiments across a number of contexts show that miRNAs produce larger changes in mRNA levels than in translation efficiency, leading the authors to suggest that mRNA destabilization is the dominant effect of miRNA repression (Eichhorn et al., 2014a; Guo, Ingolia, Weissman, & Bartel, 2010a). However, in other studies, it has been suggested that miRNAs primarily inhibit translation levels. For example, in the early zebrafish embryo, ribosome profiling and RNA-Seq show that miRNAs induce translational repression without mRNA destabilization (Bazzini, Lee, & Giraldez, 2012). Furthermore, experiments using miRNA reporters to examine the kinetics of miRNA repression suggest that translational repression precedes mRNA destabilization (Béthune, Artus-Revel, & Filipowicz, 2012; Djuranovic, Nahvi, & Green, 2012). These studies raise the question of whether translational repression is the direct mode of miRNA-driven suppression and whether mRNA destabilization is a secondary consequence. To resolve this question, it is important to genetically separate the two functions. However, despite extensive research, it is not known whether it is possible to decouple miRNA-induced translational repression and mRNA destabilization of endogenous transcripts in a cell where both occur.

The RNA-binding protein DDX6 and its yeast homolog DHH1 are DEAD box helicases that localize to P-bodies and stress granules and have been implicated in both translational repression and mRNA destabilization, suggesting that they may link these two processes (Coller & Parker, 2005; Vlad Presnyak & Coller, 2013). Tethering experiments in yeast that lack DCP2 or DCP1 demonstrate that DHH1 can repress translation upstream and independent of enhancing decapping (Carroll, Munchel, & Weis, 2011; Sweet, Kovalak, & Coller, 2012).

Tethering experiments in human cells demonstrate that DDX6 also represses translation (Kuzuoğlu-Öztürk et al., 2016a). Furthermore, DDX6 binds to components of the decapping complex, but exactly how this impacts translation and mRNA stability is unclear (Ayache et al., 2015;Nissan, Rajyaguru, She, Song, & Parker, 2010;Tritschler et al., 2009). Additionally, through interactions with the CCR4-NOT complex and the decapping complex, DDX6 is thought to be involved in miRNA mediated translational repression, but its exact role is not fully understood (Y.Chen et al., 2014a; Chu & Rana, 2006;Mathys et al., 2014a;Rouya et al., 2014a).

Here we sought to understand how mRNA stability changes are linked to translation changes during early mammalian development. It has been suggested that up to 70% of the molecular changes during mouse embryonic stem cell (ESC) differentiation are due to post-transcriptional regulation (Lu et al., 2009a). Therefore, we measured and analyzed changes in mRNA stability and translation efficiency during ESC differentiation. Surprisingly, we found that the vast majority of molecular changes during this transition are driven by transcriptional, not post-transcriptional mechanisms. However, within self-renewing ESCs there was a wide range of mRNA stabilities. These stability differences correlated with translational levels. We generated *Ddx6* KO ESCs to determine whether DDX6 links translation to stability. Unlike its yeast homolog, DDX6 did not appear to play a general role in linking the two. However, its loss did lead to the translational upregulation of miRNA targets with little associated changes in mRNA stability. The resulting cells looked phenotypically and molecularly similar to cells deficient for all miRNAs. Therefore, the loss of DDX6 is able to separate the two central functions of miRNAs: translational repression and mRNA destabilization. Furthermore, these data show miRNA induced translational repression alone can recapitulate many of the downstream consequences of miRNAs.

## RESULTS

### Transcriptional changes drive expression changes during the ESC to EpiLC transition

Previous work suggested that up to 70% of the molecular changes that occur during early ESC differentiation are due to post-transcriptional events (Lu et al., 2009b). In that work, differentiation was induced by expressing a shRNA to Nanog in ESCs grown in LIF. These conditions are associated with a heterogeneous population of cells (Ivanova et al., 2006). To revisit this question, we first turned to a reporter system and an optimized differentiation protocol that enables the homogenous differentiation of naive ESCs to formative epiblast like cells (EpiLC), which is representative of the transition from the pre- to post-implantation epiblast in vivo (A. Chen et al., n.d.;Krishnakumar et al., 2016;Parchem et al., 2014) (Figure 1A). Using this system, we characterized the changes in mRNA expression, mRNA stability, and translation that occur during the transition. RNA-Seq showed 1890 genes significantly up-regulated and 1532 genes significantly down-regulated during the ESC to EpiLC transition (Figure 1B, 1F). Known naive markers were down-regulated, while known primed markers were up-regulated during this transition (Figure 1 - figure supplement 1A) (Boroviak et al., 2015).

**Figure 1.**
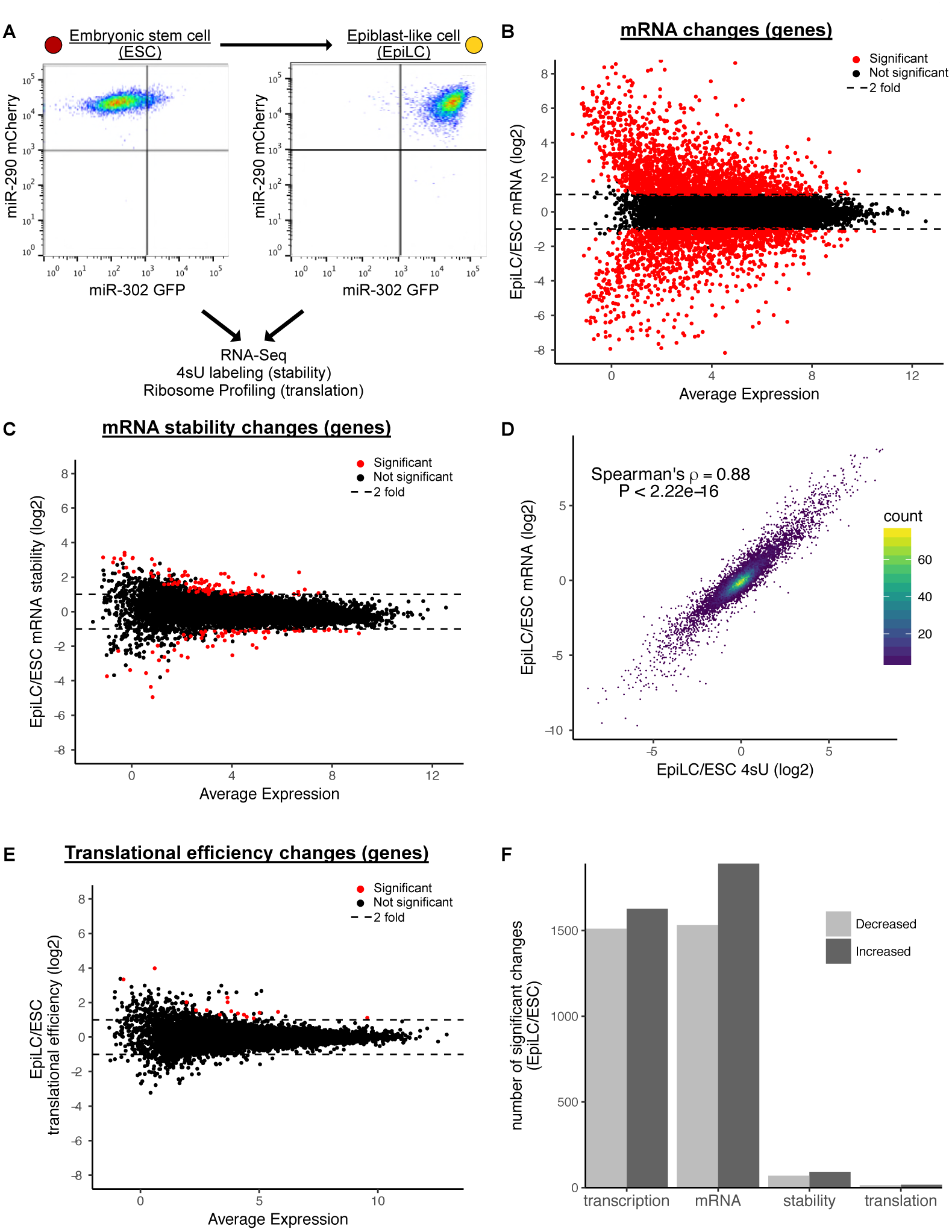
Transcriptional changes drive expression changes during the ESC to EpiLC transition. A) Flow cytometry of the transition from naïve embryonic stem cells (ESCs) (miR-302 GFP-, miR-290 mCherry+) to primed epiblast-like cells (EpiLCs) (miR-302 GFP+, miR-290 mCherry+). B) MA plot of mRNA changes during the ESC to EpiLC transition. Significant changes are shown as red dots (Adjusted P Value < 0.05 and |log2 fold change| > 1) in B, C, E. Dashed lines indicated a 2 fold change. C) MA plot of mRNA stability changes during the ESC to EpiLC transition. D) Correlation between changes in nascent transcription (4sU labeled mRNA) and changes in mRNA levels during the ESC to EpiLC transition. P value calculated with correlation significance test. E) MA plot of translational efficiency (TE) changes during the ESC to EpiLC transition. F) The number of significant increases or decreases in transcription, mRNA levels, mRNA stability, and translational efficiency during the ESC to EpiLC transition. N = 3 for each ESC and EpiLC seq experiment. See also figure 1 - figure supplement 1.

In order to directly measure changes in transcription during the ESC to EpiLC transition, we used metabolic labeling with 4-thiouridine (4sU) (Dolken et al., 2008;Rabani et al., 2011;Windhager et al., 2012). Nascent transcripts were labeled with a thirty-minute 4sU pulse, biotinylated, pulled down with streptavidin, and sequenced (4sU-Seq). Total RNA-Seq was performed in parallel and the ratio of nascent RNA to total RNA was used to calculate relative stabilities for each gene (Rabani et al., 2011). To validate these findings, a subset of genes spanning a range of stabilities were measured using an alternative method where transcription was blocked with actinomycin D and mRNA levels followed over a time course by RT-qPCR (Figure 1 - figure supplement 1B, 1C). The relative stabilities predicted by the two approaches were highly correlated. Given that the 4sU-Seq approach avoids the secondary effects associated with blocking all transcription, we used those data for genome-wide analysis (Bensaude, 2011; Lugowski, Nicholson, & Rissland, 2017). Surprisingly, the 4sU/total mRNA data showed very few changes in mRNA stability between the ESC and EpiLC states (Figures 1C, 1F). This lack of changes was not because of noise among the replicates, as biological replicates were well correlated (Figure 1 - figure supplement 1D).

The general lack of changes in mRNA stability suggested that transcription is the dominant regulator of the mRNA changes during the ESC to EpiLC transition. Indeed, fold changes in total mRNA levels correlated extremely well with fold changes in 4sU labeled nascent transcripts (Spearman’s rho 0.88; P < 2.22*10^-16) (Figure 1D). Thus, nascent transcription, not mRNA stability, underlies the mRNA changes associated with the ESC to EpiLC transition.

Next, we asked whether changes in translational levels play an important role in the ESC to EpiLC transition. To measure translational levels of all genes, we performed ribosome profiling to collect Ribosome Protected Footprints (RPFs) and matched total mRNA (Ingolia, Lareau, & Weissman, 2011). As expected, RPFs showed a strong 3 nucleotide phasing of reads that was not present in the mRNA samples, confirming the quality of the data (Figure 1 - figure supplement 1E). We calculated translation using the ratio of RPF/mRNA, also known as translational efficiency (Ingolia et al., 2011). Global analysis showed very few changes in translational efficiencies between the ESC and EpiLC states (Figures 1E, 1F). Biological replicates were well correlated showing that the overall lack of changes is not due to noise between the replicates (Figure 1 - figure supplement 1F). Therefore, like mRNA stability, there are few changes in translation levels, independent of mRNA levels, in early ESC differentiation.

### There is a wide range of RNA stabilities which are positively correlated with translation level in ESCs

Although there were minimal changes in mRNA stability during the ESC to EpiLC transition, there was a wide range of mRNA stabilities within ESCs. For example, between the 25th and 75th percentile of mRNA stability, there was a 3.2 fold difference in stability and between the top and bottom 1% of mRNA stability there was over a 64 fold difference (Figure 2A). To identify features that explain the range of mRNA stabilities observed, we performed multiple linear regression taking into account the following features that previous studies implicated in affecting mRNA stability: 3’ UTR length, 5’ UTR length, CDS length, 3’ UTR GC content, 5’ UTR GC content, CDS GC content, AU rich elements (ARE), miRNA binding sites, number of exons in the transcript, and upstream ORFs (Chan et al., 2017; Cheng, Maier, Avsec, Rus, & Gagneur, 2017;Sharova et al., 2009). Combined, these features explained 25% of the variation in mRNA stability. To identify which features had the greatest impact on stability, we analyzed the correlation between each individual feature and mRNA stability (Figure 2B). 3’ UTR length had the greatest impact and was negatively correlated with mRNA stability (Spearman’s rho-0.3; P < 2.22*10^-16) (Figure 2C). To validate the impact of 3’ UTRs on mRNA stability, we used a dual reporter system that contains a control GFP for normalization and a RFP with a cloned endogenous 3’ UTR from 12 representative genes (Figure 2D) (Chaudhury et al., 2014). Flow cytometry analysis of cells expressing the reporter showed that the RFP/GFP ratio correlated well with the mRNA stability of the matching endogenous genes as measured by 4sU-Seq (Figure 2D).

**Figure 2.**
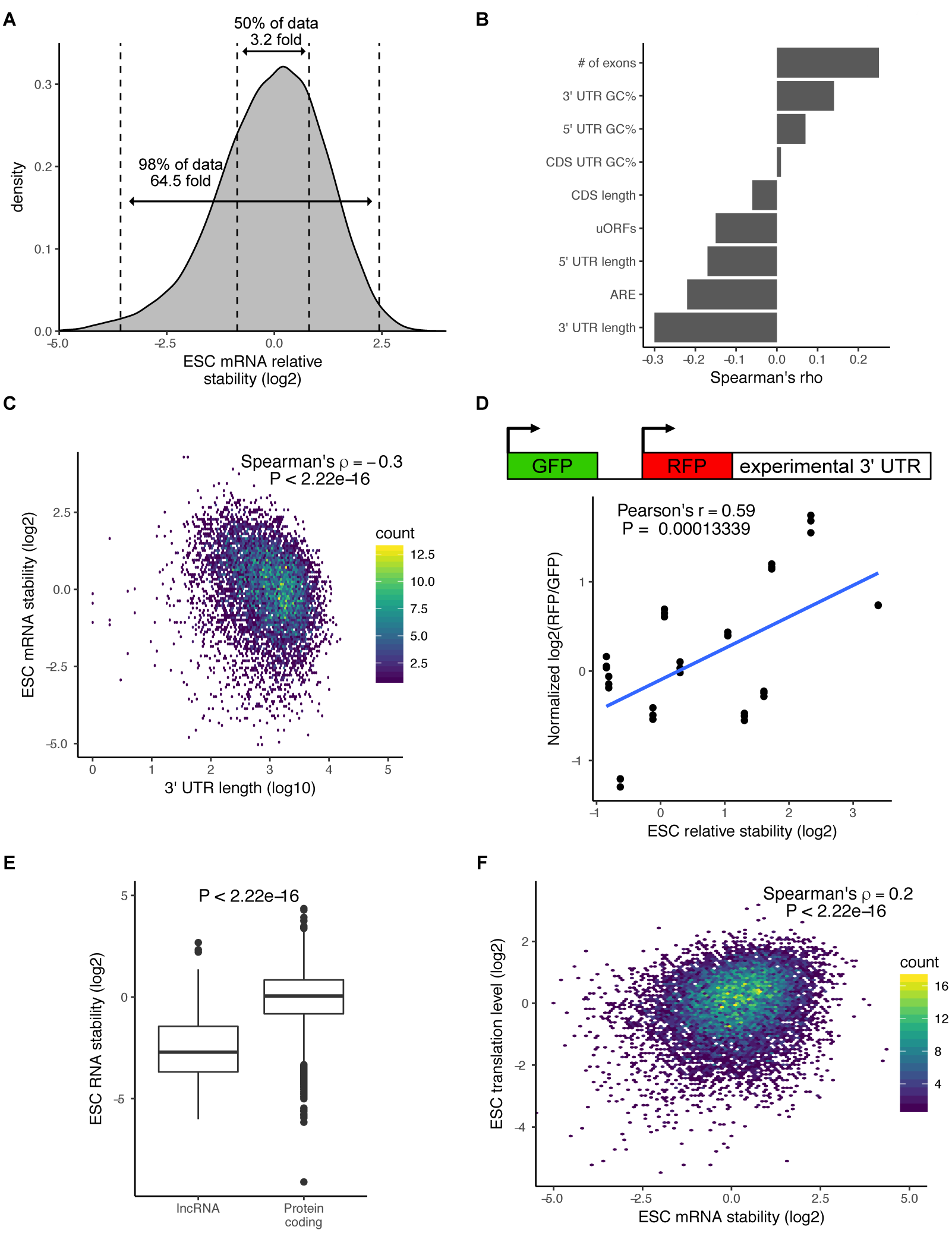
There is a wide range of RNA stabilities which are positively correlated with translation level in ESCs. A) The distribution of mRNA stabilities in ESCs. Dashed lines divide bottom 1%, middle 50%, and top 1% of the data. B) The correlation between sequence features and mRNA stability in ESCs. uORFs (upstream open reading frames), ARE (AU Rich Elements). C) The correlation between 3’ UTR length (log10) and mRNA stability in ESCs. D) (Top) schematic of dual reporter system to test endogenous 3’ UTRs. (Bottom) Normalized median RFP/GFP ratios versus mRNA stability for endogenous genes as measured by 4sU-Seq. Clusters of dots indicate an endogenous 3’ UTR, individual dots within a cluster represent biological replicates (n = 3). E) RNA stability of long non-coding RNAs (lncRNAs) compared to protein coding RNAs. P value was calculated using the Mann–Whitney test. F) Comparison between mRNA stability and translation level (high polysome / monosome ratio) in ESCs. P value calculated with correlation significance test. n = 3. See also figure 2 - figure supplement 1.

MiRNAs are one regulatory factor that bind to the 3’ UTR of target mRNAs and recruit a complex of proteins that then destabilize the transcripts (Fabian & Sonenberg, 2012; Jonas & Izaurralde, 2015). In ESCs the embryonic stem cell-enriched cell cycle (ESCC) family of miRNAs represent a predominant fraction of all miRNAs in ESCs (Greve, Judson, & Blelloch, 2013a; Houbaviy, Murray, & Sharp, 2003a;Marson et al., 2008a). As expected, ESCC miRNA targets as a group were significantly less stable than all genes (P < 2.22*10^-16, Mann-Whitney test) (Figure 2 - figure supplement 1A). However, a large number of ESCC targets were still in the top 50% of the most stable genes (Figure 2 - figure supplement 1A). This data suggests that, while miRNAs are strong destabilizers, they can only explain a small portion of the large range of mRNA stabilities seen in the cells.

Interestingly, analysis of the 4sU-Seq data showed that long non-coding RNAs (lncRNAs) were significantly less stable than protein coding genes (P < 2.22*10^-16, Mann-Whitney test) (Figure 2E). This result suggested a strong role for translation in regulating RNA stability in ESCs. To further test this hypothesis, we performed polysome profiling (Arava et al., 2003). While both ribosome profiling and polysome profiling measure global levels of translation, polysome profiling enables the measurement of relative elongation time versus initiation time by comparing the fraction of transcripts bound by multiple ribosomes (polysome fraction) versus a single ribosome (monosome fraction) (Heyer & Moore, 2016). Thus, it can be a more sensitive measure of translation regulation. We isolated monosome fractions, low polysome fractions containing 2-4 ribosomes, and high polysome fractions containing 4+ ribosomes and performed RNA-Seq (Figure 2 - figure supplement 1B). To avoid confusion with the translation efficiency metric measured by ribosome profiling, we refer to the ratio of high polysome/monosome as translation levels. As expected, protein coding genes had a much higher translation level compared to lncRNAs (P < 2.22*10^-16, Mann-Whitney test) (Figure 2 - figure supplement 1C). We next compared the polysome profiling data to the mRNA stability data. There was a highly significant correlation between mRNA stability and translation levels across all genes (Spearman’s rho 0.2; P < 2.22*10^-16) (Figure 2F). These findings suggested a direct link between translation level and mRNA stability.

Recent reports suggest that differential codon usage is a central mechanism in linking protein translation to mRNA stability (Bazzini et al., 2016;Chan et al., 2017;Cheng et al., 2017; Mishima & Tomari, 2016; Vladimir Presnyak et al., 2015). Therefore, we considered the possibility that codon optimality is a driving force in the wide range of mRNA stabilities. Analysis of the frequency that each codon appears in a transcript uncovered small differences in codon usage frequency when comparing the top and bottom 20% of mRNA stabilities (Figure 2 - figure supplement 1D). Therefore, codon optimality may in part explain the link between translation levels and mRNA stability.

### DDX6 regulates proliferation and morphology of ESCs

In yeast, the protein DHH1 has been shown to link translation to mRNA stability through codon optimality (Radhakrishnan et al., 2016). The mammalian homolog of DHH1, DDX6, has been shown to associate with both the mRNA decapping and deadenylation complex, also consistent with a potential link between mRNA stability and translation (Y. Chen et al., 2014b;Mathys et al., 2014b;Rouya et al., 2014b). Therefore, we next asked whether DDX6 may provide a mechanistic link for the relationship between translation and mRNA stability in ESCs. To investigate the function of DDX6 in ESCs, we produced *Ddx6* knockout (*Ddx6* KO) clones using CRISPR-Cas9. Sanger sequencing confirmed a single nucleotide insertion in one clone and a large deletion in a second clone, both of which produce a premature stop (Figure 3 - figure supplement 1A). Western blot confirmed the absence of DDX6 protein in both clones (Figure 3A). We repeated the 4sU-Seq and polysome profiling in *Ddx6* KO and matched wild type cells to measure changes in mRNA stability and translation. 4sU-Seq and total RNA-Seq showed that while there was a minimal reduction of nascent *Ddx6* mRNA, there was a drastic loss of mature *Ddx6* mRNA in the *Ddx6* KO cells (Figure 3B). This destabilization is consistent with nonsense mediated decay and further validates the 4sU-Seq assay for assessing changes in mRNA stability.

The loss of DDX6 had little impact on the expression of pluripotency markers (Figure 3C). However, there were striking morphological changes in the cells (Figure 3D). Unlike wild type ESCs which form tight domed colonies, *Ddx6* KO cells grew in a jagged, dispersed monolayer (Figure 3D). DDX6 localized to discrete punctate in the wild type cells consistent with P-body localization, as previously reported (Figure 3E) (Ernoult-Lange et al., 2012;Hubstenberger et al., 2017; Minshall, Kress, Weil, & Standart, 2009; Vlad Presnyak & Coller, 2013). Interestingly, the loss of DDX6 resulted in an abnormal distribution of the P-body marker DCP1a, suggesting a role for DDX6 P-body formation in ESCs (Figure 3F). DDX6 loss also led to a reduction in proliferation in self-renewal culture conditions (Figure 3G). Together, these data show an important role for DDX6 in retaining normal cell morphology and proliferation.

**Figure 3.**
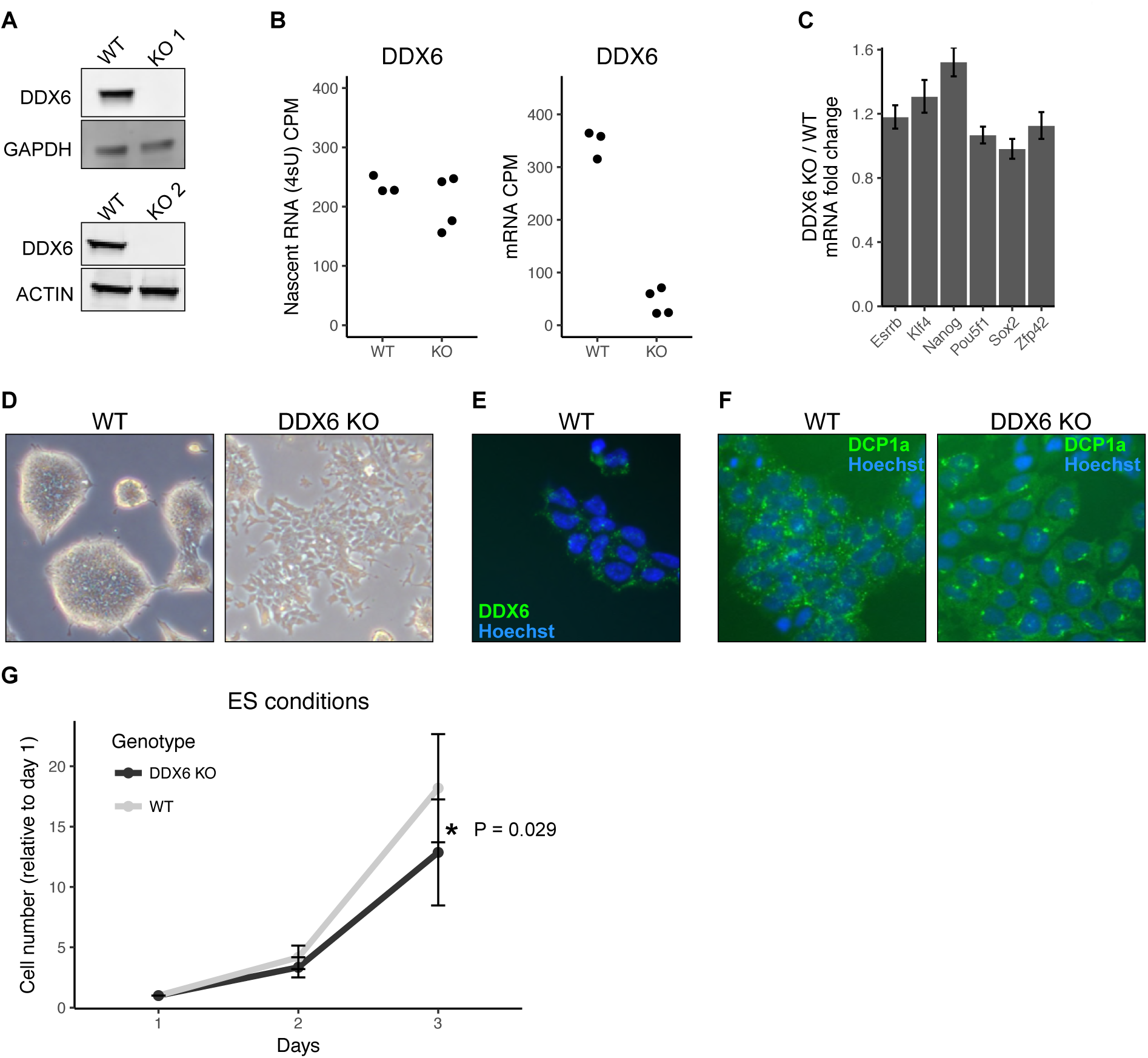
DDX6 regulates proliferation and morphology of ESCs. A) Western blot of DDX6 in two DDX6 knockout lines. GAPDH and ACTIN were used as loading controls. B) *Ddx6* counts per million (CPM) in nascent mRNA (4sU) or mRNA in wild type (WT) and *Ddx6* KO cells. n = 3 for wild type, n = 4 for *Ddx6* KO (2 replicates of each *Ddx6* KO line) C) Expression of pluripotency genes in *Ddx6* KO ESCs based on RNA-Seq. Error bars represent 95% confidence interval. D) Brightfield images of wild type and *Ddx6* KO ESCs. Images taken at 20X. E) DDX6 staining in wild type ESCs. Images taken at 20X. F) P-body staining against DCP1a in wild type and *Ddx6* KO ESCs. Images taken at 20X. G) Growth curves of wild type and *Ddx6* KO ESCs in ESC maintenance conditions (LIF/2i). n = 6 for wild type cells, n = 12 for *Ddx6* KO (6 replicates of each *Ddx6* KO line). * indicates P < 0.05 using a t-test, error bars are standard deviation. See also figure 3 - figure supplement 1.

### DDX6 separates miRNA-induced translation repression from RNA degradation

Since yeast DHH1 destabilizes lowly translated transcripts enriched in non-optimal codons, we expected that lowly translated genes might be stabilized in *Ddx6* KO ESCs (Radhakrishnan et al., 2016). However, there was minimal correlation between mRNA stability changes in *Ddx6* KO ESCs and wild type translation levels (Spearman’s rho-0.11; P < 2.22*10^-16) (Figure 4 - figure supplement 1A). The stabilized transcripts were not specifically enriched within the lowly translated transcripts and instead they occurred across all levels of translation. These data suggested that DDX6 does not link mRNA stability with translation across all genes. Next, we defined a set of codons as suboptimal based on their enrichment in unstable genes in wild type ESCs and asked whether they are enriched among genes that are stabilized in *Ddx6* KO cells (Figure 2 - figure supplement 1D). There was no enrichment (Figure 4 - figure supplement 1B). Further comparing changes in median codon frequency in stable versus unstable transcripts in wild type cells with changes in median codon frequency in stabilized versus unstabilized transcripts in *Ddx6* KO cells showed no correlation (Figure 4 - figure supplement 1C). Species specific tRNA adaptation index (sTAI) provides an alternative metric of codon optimality. The sTAI metric takes into account tRNA copy number and a tRNA’s ability to wobble base pair with different codons (Radhakrishnan et al., 2016; Sabi & Tuller, 2014). We calculated sTAI values for mouse and asked if they could predict changes in transcript stability associated with DDX6 loss. In contrast to the yeast homolog, transcripts stabilized upon DDX6 loss did not correlate with low sTAI values (Figure 4 - figure supplement 1D). These data show that unlike yeast DHH1, the primary function of mammalian DDX6 is not to link codon optimality with transcript stability.

Several aspects of the *Ddx6* KO phenotype, including the cell morphology changes and growth defects, resemble the phenotype of *Dgcr8* KO cells (Wang, Medvid, Melton, Jaenisch, & Blelloch, 2007). DGCR8 is essential for miRNA biogenesis and *Dgcr8* KO ESCs lack all miRNAs (Wang et al., 2007). DDX6 has been implicated as an effector of miRNA activity (Y.Chen et al., 2014b; Chu & Rana, 2006;Mathys et al., 2014b;Rouya et al., 2014b). Therefore, we next asked whether *Ddx6* KO cells have similar downstream molecular consequences as *Dgcr8* KO cells. To directly compare the two, we performed 4sU-Seq and polysome profiling in *Dgcr8* KO ESCs and analyzed the data in parallel with that of the *Ddx6* KO ESCs.

The embryonic stem cell enriched cell cycle (ESCC) family of miRNAs represent a predominant fraction of all miRNAs in ESCs (Greve, Judson, & Blelloch, 2013b; Houbaviy, Murray, & Sharp, 2003b;Marson et al., 2008b; Melton, Judson, & Blelloch, 2010;Wang et al., 2008). They share the “AAGUGC” seed sequence and thus have common downstream targets. Furthermore, re-introduction of a single member of the ESCC family of miRNAs can revert *Dgcr8* KO cells to a molecular phenotype highly similar to wild type ESCs (Gambardella et al., 2017;Melton et al., 2010;Wang et al., 2008). Therefore, we chose to focus on the consequence of DGCR8 loss and DDX6 loss on these targets. As expected, the ESCC targets are stabilized relative to all genes in the *Dgcr8* KO cells (Figure 4A). However, these same targets showed little change in mRNA stability in the *Ddx6* KO cells (Figure 4B). Therefore, DDX6 does not appear to play a major role in transcript destabilization downstream of miRNAs.

**Figure 4.**
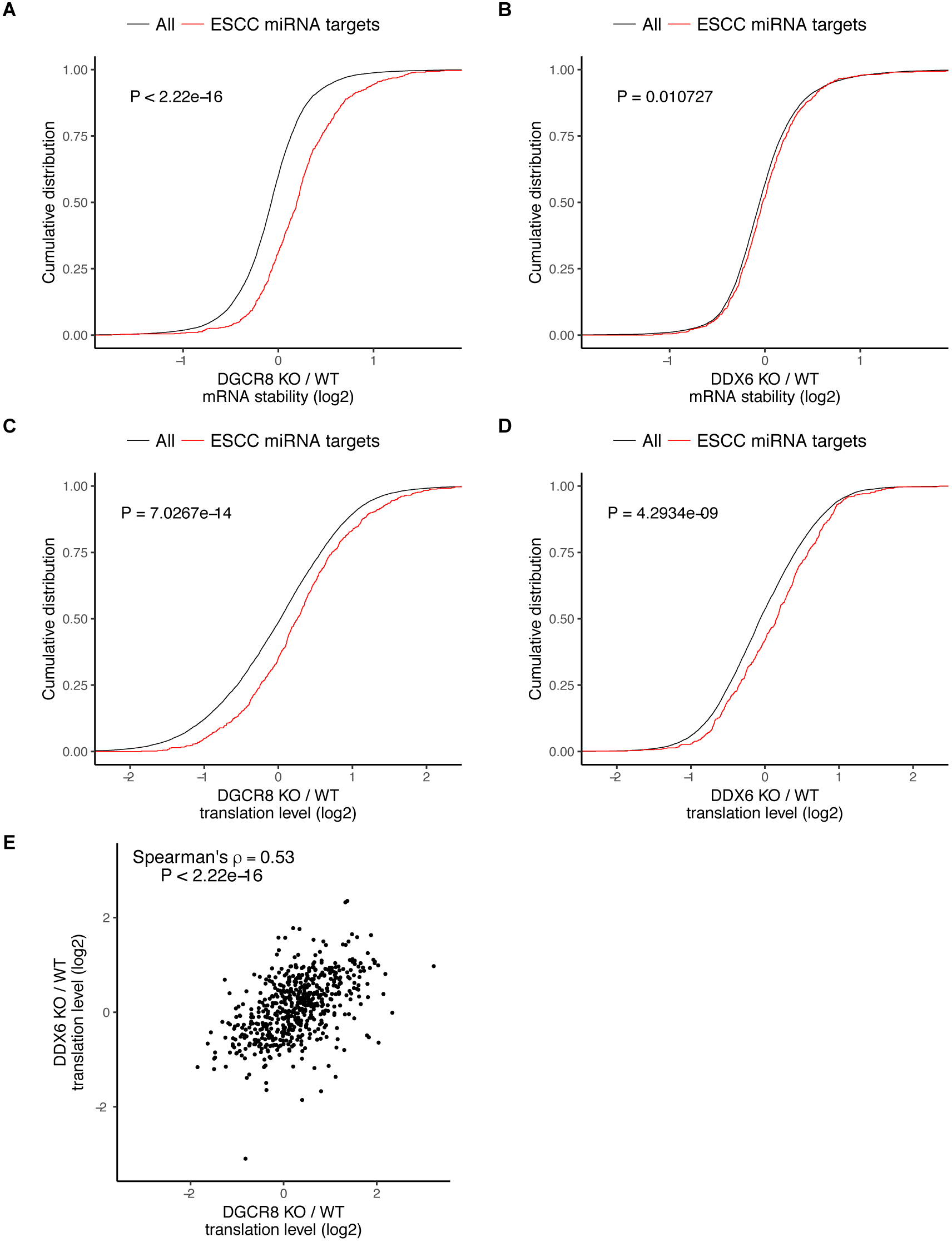
DDX6 separates miRNA-induced translation repression from RNA degradation. A-D) mRNA stability or translation level changes in ESCC miRNA targets versus all mRNAs. P value calculated with Mann-Whitney test. A/B) mRNA stability changes in *Dgcr8* KO (A) or *Ddx6* KO (B) cells. n = 3 for wild type, n = 4 for *Ddx6* KO (2 replicates of each *Ddx6* KO line), n = 3 for *Dgcr8* KO. C/D) Translation level changes in *Dgcr8* KO (C) or *Ddx6* KO (D) cells. n = 3 for each genotype. E) Translation level changes of individual ESCC miRNA targets in *Dgcr8* KO and *Ddx6* KO ESCs. See also figure 4 - figure supplement 1.

The loss of DGCR8 also resulted in an increase in translation of ESCC miRNA targets independent of its effect on stability, consistent with miRNAs both inhibiting translation and destabilizing transcripts (Figure 4C). In contrast to the stability data, the loss of DDX6 had a similar impact as the loss of DGCR8 on the translation of ESCC targets (Figure 4D). Indeed, *Dgcr8* KO and *Ddx6* KO affected the translation of individual targets to a similar extent (Figure 4E). These data show that DDX6 is an essential effector for miRNA driven translational repression, but not mRNA stabilization. As such, DDX6 separates the two main functions of miRNAs showing that miRNA driven translational repression and transcript destabilization are not dependent on one another.

### Translational repression alone underlies many of the downstream molecular changes associated with miRNA loss

Whether translational repression or mRNA destabilization is the predominant effect of miRNAs is controversial as it is difficult to separate the two (Iwakawa & Tomari, 2015; Jonas & Izaurralde, 2015). Given that the *Ddx6* KO cells retained mRNA destabilization, while losing translational repression of miRNA targets, we asked how well derepression of translation matches the downstream consequences of losing all miRNAs. Since the *Ddx6* KO and *Dgcr8* KO cells have partially overlapping phenotypes, we compared global changes in mRNA stability, mRNA levels, and translation levels. Strikingly, while there was little correlation in changes in mRNA stability, changes in both mRNA and translation levels were well correlated (Figure 5). The correlation in mRNA changes is likely due to transcriptional secondary effects due to losing translation repression of miRNA targets in *Ddx6* KO and *Dgcr8* KO cells. These data show that translational repression alone can explain much of a miRNA’s function in ESCs.

**Figure 5.**
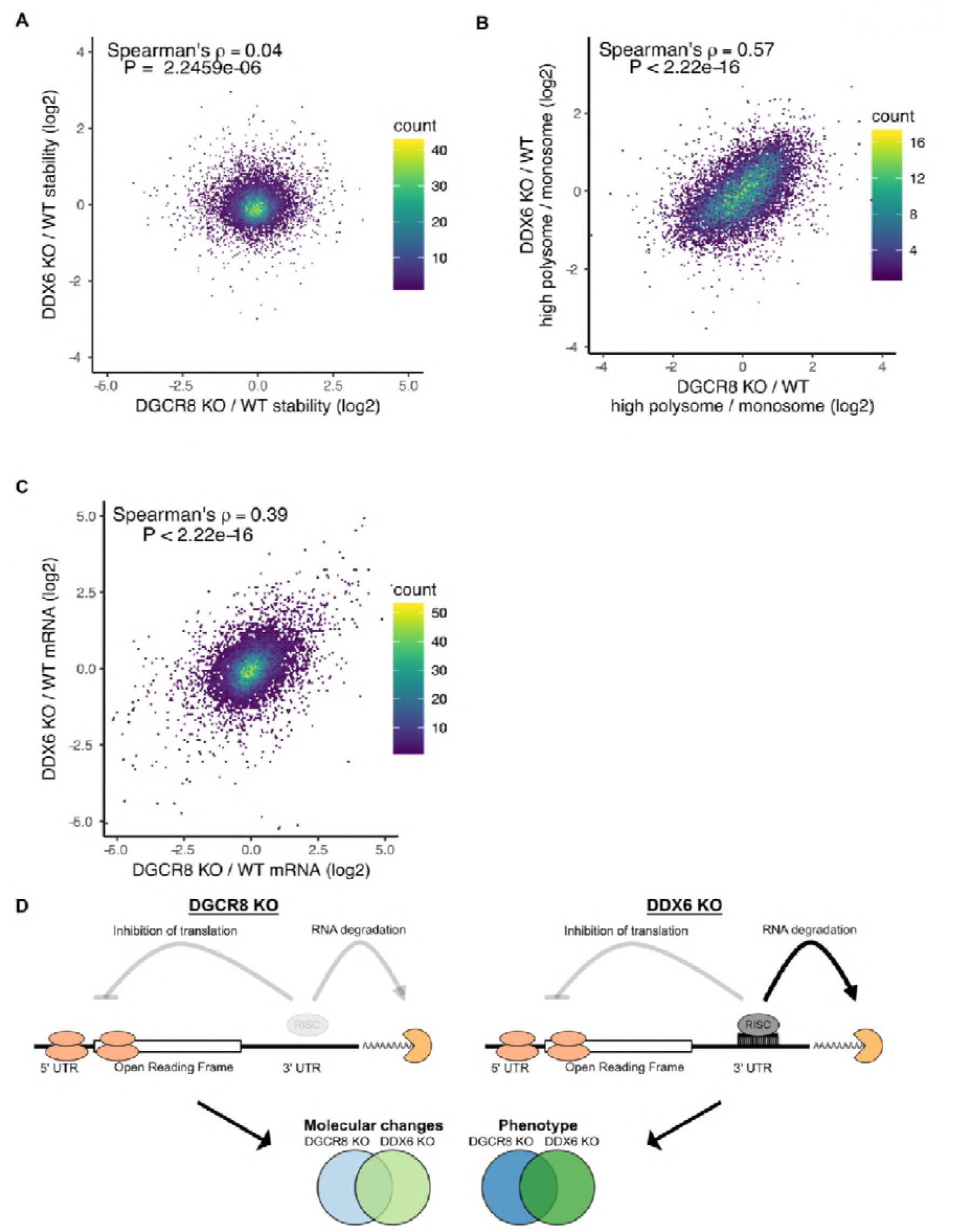
Translational repression alone underlies many of the downstream molecular changes associated with miRNA loss. A) Comparison between mRNA stability changes in *Dgcr8* KO versus *Ddx6* KO cells. n = 3 for wild type, n = 4 for *Ddx6* KO (2 replicates of each *Ddx6* KO line), n = 3 for *Dgcr8* KO. B) Comparison between translation level changes in *Dgcr8* KO versus *Ddx6* KO cells. n = 3 for each genotype. C) Comparison between mRNA changes in *Dgcr8* KO versus *Ddx6* KO cells. P value calculated with correlation significance test. D) Summary schematic comparing *Dgcr8* KO cells to *Ddx6* KO cells.

## DISCUSSION

In this study we sought to uncover how mRNA stability and translation are regulated within the ESC state and during differentiation. Using RNA-Seq, metabolic labeling (4sU-Seq), and ribosome profiling, we found that most changes during ESC differentiation are driven at the level of transcription. This finding contrasts Lemischka and colleagues’ conclusion that post-transcriptional changes underlie many changes in protein levels during ESC differentiation (Lu et al., 2009a). These differences are likely explained by the different approaches used and the fact that Lemischka and colleagues focused their analysis on nuclear protein changes. Discordant changes between mRNA expression and nuclear protein levels could reflect changes in translation levels, protein stability, or protein localization (Liu, Beyer, & Aebersold, 2016). We measured translation levels, but not post-translational events; therefore, it remains plausible that protein degradation rates or protein localization are dynamic during ESC differentiation.

We found a positive correlation between mRNA stability and translation levels in ESCs, similar to what other groups have observed recently in yeast (Chan et al., 2017; Heyer & Moore, 2016; Vladimir Presnyak et al., 2015). A number of mechanisms have been proposed. One is that ribosomes sterically hinder the degradation machinery from accessing the transcript. In support of this model, RNA-Seq of the 5’ end of decapped RNA degradation intermediates shows a three nucleotide periodicity consistent with exonucleases running into the ribosome on a final round of translation (Pelechano, Wei, & Steinmetz, 2015). Alternatively, some RNA-binding proteins may sense slowly translating transcripts and accelerate their degradation as recently described for DHH1 (Radhakrishnan et al., 2016). However, in our data the loss of the mammalian homolog of DHH1, DDX6, did not appear to link low levels of translation with low mRNA stability.

There has been extensive debate about whether miRNAs primarily inhibit translation or induce destabilization of their target transcripts (Iwakawa & Tomari, 2015; Jonas & Izaurralde, 2015). By analyzing stability changes and translation changes in *Dgcr8* KO ESCs, which lack all mature miRNAs, we observe that miRNAs both inhibit translation and induce destabilization within ESCs. However, upon loss of DDX6, miRNA targets are derepressed at the translational level without affecting miRNA induced destabilization. It has been suggested that translational repression of miRNA targets is the cause of mRNA destabilization or is at least a prerequisite (Radhakrishnan & Green, 2016). However, the *Ddx6* KO cells demonstrate that mRNA destabilization can occur without translational repression in a context where both forms of repression normally occur. Future studies will likely identify factors that can decouple translational repression and mRNA destabilization in the other direction so that miRNA targets are translationally repressed without inducing mRNA destabilization. This situation has been observed in the early zebrafish embryo, but the mechanism underlying the phenomenon remains unclear (Bazzini et al., 2012).

Our data supports a key role for DDX6 in the translational repression of miRNA targets. However, it is not fully understood how DDX6 represses translation of these targets. Experiments using DDX6 tethered to different reporters suggest that DDX6 suppresses translational initiation (Kuzuoğlu-Öztürk et al., 2016b). It was recently shown that DDX6 interacts with 4E-T, which competes with eIF4G for binding to the translation initiation factor eIF4E and leads to translational repression (Kamenska et al., 2016;Ozgur et al., 2015). Additionally, mutations in the FDF binding domain of DDX6 prevents interaction with 4E-T and decapping proteins and prevents translational repression of a reporter (Kuzuoğlu-Öztürk et al., 2016c). However, depletion of 4E-T only partially alleviates DDX6 mediated translational repression (Kamenska et al., 2014;Kuzuoğlu-Öztürk et al., 2016c). Therefore, DDX6 likely interacts with additional unknown factors to inhibit translation initiation. Our data shows that the loss of DDX6 results in increased translation of miRNA targets to a similar level as the loss of all miRNAs, suggesting that DDX6 serves as a key link between the proteins that repress translation and the rest of the RNA induced silencing complex.

Surprisingly, we found that *Dgcr8* KO and *Ddx6* KO have similar overall downstream molecular consequences, even though the former leads to both stabilization and translational derepression of miRNA targets, while the later only influences translation. Furthermore, *Ddx6* KO and *Dgcr8* KO cells both display morphology defects and proliferation defects. It has been argued that mRNA changes are the dominant effect of miRNAs, since miRNA induced changes in mRNA levels are often larger than changes in translation efficiency (Eichhorn et al., 2014b; Guo, Ingolia, Weissman, & Bartel, 2010b). However, our data suggests that while miRNAs often have a significant effect on mRNA stability, their impact on translational alone can recapitulate a large portion of the downstream molecular and phenotypic effects associated with miRNA loss.

## MATERIALS AND METHODS

### Gene accession

The accession number for the sequencing data reported in this paper is GEO: GSE112767.

### 4sU-Sequencing

Samples were labeled with 500uM 4-thiouridine (4sU) (Sigma) for 30 minutes then extracted with TRIzol (Invitrogen) and split into two groups. rRNA was depleted from Total RNA using the Ribo-Zero Gold kit (Illumina). 80 ug of RNA was biotinylated according to the following protocol Radle et al. (Rädle et al., 2013). Biotinylated 4sU RNA was isolated and washed using M-270 Streptavidin Dynabeads (Invitrogen), eluted with 100mM DTT, and cleaned up with Rneasy minelute columns (Qiagen).

Libraries were generated with the KAPA Stranded RNA-Seq or Stranded HyperPrep library prep kit (Kapa) and sequenced with single-end 50bp reads. Additional rounds of *Ddx6* KO and matched wild type 4sU samples were sequenced with paired-end reads and counts were merged with single end reads.

### Cell culture and differentiation

ESCs were grown in Knockout DMEM (Invitrogen) supplemented with 15% Fetal Bovine Serum, LIF and 2i (PD0325901 and CHIR99021). In order to generate EpiLCs, 400,000 ESCs were plated in a 15cm plate; 24 hours later LIF/2i media was removed, cells were washed with PBS, and EpiLCs were collected ~ 56 hours later (Krishnakumar et al., 2016). Cells were tested to be free of mycoplasma.

### Quant Seq

QuantSeq 3’ end counting was used for polysome profiling samples as well as matched wild type, *Ddx6* KO, and *Dgcr8* KO mRNA samples (Figure 4F). RNA was isolated using Rneasy Micro kits (Qiagen). RNA-Seq libraries were generated using the QuantSeq 3’ FWD kit (Lexogen) and sequenced with single-end 50bp reads.

### Ribosome profiling

ESCs and EpiLCs were grown as above. Ribosome profiling libraries were generated using the TruSeq Ribosome Profiling kit (Illumina) and sequenced with single-end 50bp reads. 3 nucleotide periodicity of ribosome profiling reads was checked using RiboTaper (Calviello et al., 2016). Adapters were trimmed using cutadapt version 1.14 with the following settings:--minimum-length 26 –maximum-length 32 for the ribosome protected fragments or –minimum-length 32 for the total RNA. Adapater sequence used for trimming: AGATCGGAAGAGCACACGTCT. Reads were mapped with STAR version 2.5.3a to the mm10/Gencode M14 genome with the following settings:--outFilterMultimapNmax 1 – outFilterMismatchNoverReadLmax 0.05 –seedSearchStartLmax 13 –winAnchorMultimapNmax 200.

### Polysome Profiling

Two plates of 6 million ESCs were seeded in a 15 cm plate 48 hours prior to collection. Cells were incubated with 100 ug/ml cycloheximide (Sigma) for 2 minutes and then moved to ice. Cells were washed and scraped in PBS with cycloheximide, spun down, and then lysed. Lysate was loaded onto a 10-50% sucrose gradient and centrifuged at 35,000 RPM for 3 hours. Gradients were collected on a gradient station (Biocomp). For each sample, the monosome, low polysome (2-4 ribosomes), and high polysome (4+ ribosomes were collected). RNA was extracted from gradient fractions with TRIzol LS (Invitrogen) and concentrated with the Zymo Clean and Concentrator-5 kit (Zymo) prior to library preparation with the QuantSeq 3’ FWD kit (Lexogen).

### Western blot

Cells were collected in RIPA buffer with Protease Inhibitor Cocktail (Roche). Protein was run on a 4-15% gel (Bio-Rad) then transferred onto a PVDF membrane. Membranes were blocked with Odyssey blocking buffer, blotted with primary and secondary antibodies, and then imaged on the Odyssey imaging system (LI-COR). Antibodies: DDX6 1:1000 (A300-460A-T), GAPDH 1:1000 (SC 25778), ACTIN 1:1000 (A4700).

### Actinomycin D RT-qPCR

Cells were treated with 5 ug/ml Actinomycin D. 0, 2, 4, 6, 8, and 12 hours after treatment, RNA was collected in TRIzol (Invitrogen). Reverse transcription was performed with the Maxima first strand synthesis kit (Thermo Scientific). qPCR was then performed with the SensiFAST SYBR Hi-ROX kit (Bioline) on an ABI 7900HT 384-well PCR machine. Each sample was normalized to 18S rRNA and its 0 hour time point.

### Differentiation RT-qPCR

Cells were differentiated to an EpiLC state as described above, scaled down to a 6 well dish. Samples were collected in TRIzol (Invitrogen). Reverse transcription was performed with the Maxima first strand synthesis kit (Thermo Scientific). qPCR was then performed with the SensiFAST SYBR Hi-ROX kit (Bioline) on an ABI 7900HT 384-well PCR machine. Each sample was normalized to Gapdh and wild type ESCs in LIF/2i conditions.

### BAX/BAK RT-qPCR

Cells were collected in TRIzol (Invitrogen) and RNA was isolated using the Directzol miniprep kit (Zymo). qPCR was then performed with the SensiFAST SYBR Hi-ROX kit (Bioline) on an ABI 7900HT 384-well PCR machine. Each sample was normalized to Gapdh and the parental BAX/BAK cell line.

### Cell count

50,000 cells were plated in multiple wells of a 6 well on day 0. On day 1, 2, and 3 cells were trypsinized and counted with a TC20 (Bio-rad). Day 2 and 3 counts were normalized to the day 1 count.

### Imaging

Cells were fixed with 4% PFA 10 minutes at room temperature. Cells were blocked with 2% BSA and 1% goat serum in PBST. Cells were incubated with primary antibody for 1 hour at room temperature (Dcp1 abcam (ab47811) antibody 1:800 or DDX6 A300-460A) antibody 1:250). Cells were incubated with goat 488 secondary for 1 hour at room temperature. Cells were then imaged on a Leica inverted fluorescence microscope.

### Generation of *Ddx6* KO ESCs

*Ddx6* KO lines were generated using the protocol from Ran et al. (Ran et al., 2013). A guide RNA (CATGTGGTGATCGCTACCCC) was cloned into PX458, transfected into ESCs using Fugene 6, and then GFP positive cells were sorted at clonal density. Clones were genotyped with the following primers (Fwd: CATTGCCCAGATTGAAGACA and Rvs: TCCTGACTGGCCTGAAACTT) and verified by western blot. Two different knockout clones were picked and used for all subsequent analysis.

### Species specific tRNA adaptation index calculation

For each gene, the CDS region from the Gencode M14 annotation was used. Species specific tRNA adaptation index (sTAI) values for each gene were calculated with stAIcalc (Sabi, Volvovitch Daniel, & Tuller, 2017).

### Calculation of codon usage

For each gene, the APPRIS principle isoform was used to calculate codon usage frequency. To analyze differences in codon usage between stable and unstable genes, codon usage frequency was calculated for genes in the top 20% (stable) and bottom 20% (unstable) in terms of wild type mRNA stability. For codon usage frequency for mRNA stability changes in *Ddx6* KO cells, we first filtered for genes in the bottom 20% of wild type stability as defined above. Within those genes, we took the top 20% (top) and bottom 20% (bottom) of mRNA stability changes in *Ddx6* KO ESCs and calculated codon usage frequency within each group. For the comparison between codon usage frequency in wild type versus *Ddx6* KO, we took the median codon usage frequency in stable – the median codon usage frequency in unstable for each codon and compared it to the *Ddx6* KO median codon usage frequency in the bottom group – median codon usage frequency in the top group, using groups as defined above.

### Analysis software

For all samples, adapters were trimmed with Cutadapt version 1.14 with the following options: - m 20-a “A{18}”-a “T{18}”-a AGATCGGAAGAGCACACGTCTGAACTCCAGTCAC. Reads were mapped with STAR version 2.5.3a to the mm10/Gencode M14 genome with the following settings: --outFilterMultimapNmax 1 –outFilterMismatchNoverReadLmax 0.05 – seedSearchStartLmax 25 –winAnchorMultimapNmax 100. Reads were counted with featureCounts version 1.5.3 using the Gencode M14 annotation with rRNA annotations removed with the following settings:-s. Differential expression was carried out with limma version 3.32.10 and R version 3.4.2. Genes with a low number of reads were filtered out: a gene must have at least 3 counts per million across at least 3 replicates to be included for differential expression. For samples with multiple comparisons, a linear model was used for each condition in limma taking into account assay type (e.g. 4sU versus total RNA) and cell type (e.g. KO versus wild type); significant changes in stability or translation are based on the interaction term. All downstream analysis was performed in R version 3.4.2 and plotted with ggplot2.

### Polysome Profiling Analysis

RNA-Seq from the monosome, low polysome (2-4 ribosomes), and high polysome (4+ ribosomes were collected) fractions was mapped as above. Translation level was defined as the ratio of the high polysome counts divided by the monosome counts. For KO versus wild type analysis, a linear model was used for each condition in limma and significant changes in translation are based on the interaction term.

### 4sU-Seq Analysis

By measuring transcription rate and steady state mRNA levels, it is possible to infer the relative degradation rate (Rabani et al., 2011). It is assumed that across the population of cells there is no change in mRNA levels over time for a given state. Therefore, changes mRNA levels can be modeled by their production rate *α* and degradation rate *β*.

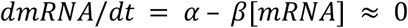

Solving for this equation, degradation rates can be calculated using a production rate (in this case nascent RNA transcription as measured by 4sU incorporation) and the concentration of total mRNA in the cell (as measured by total RNA-Seq).

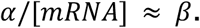

For KO versus wild type analysis, a linear model was used for each condition in limma and significant changes in translation are based on the interaction term.

### Analysis of features regulating RNA stability

For each gene with multiple isoforms, the APPRIS principle isoform was used. APPRIS data were downloaded on 10/30/2017. Log10(feature lengths), GC %, and number of exons were calculated in R version 3.4.2. Upstream open reading frames were defined as the number of ATG sequences in the 5’ UTR. AU rich elements were defined as the number of UAUUUAU sequences in the 3’ UTR. miRNA sites were defined as below. Each of these features and mRNA stability were used in a multiple linear regression using the lm function in R version 3.4.2. Additionally, the Spearman correlation was calculated between each feature and mRNA stability.

### microRNA targets

Conserved microRNA targets were downloaded from Targetscan mouse release 7.1. This list was filtered for genes that are targeted by the miR-291-3p/294-3p/295-3p/302-3p family yielding 765 target genes.

### 3’ UTR analysis

For each gene, the APPRIS principle isoform was used to calculate log10 (3’ UTR length). Log10(3’ UTR length) was then compared to log2 relative mRNA stability.

### 3’ UTR reporters

Endogenous 3’ UTRs from the following genes were amplified from ESC cDNA: ENSMUSG00000021583, ENSMUSG00000029580, ENSMUSG00000043716, ENSMUSG00000010342, ENSMUSG00000021665, ENSMUSG00000024406, ENSMUSG00000052911, ENSMUSG00000058056, ENSMUSG00000020105, ENSMUSG00000026003, ENSMUSG00000020038, ENSMUSG00000025521, ENSMUSG00000031503. Genes were cloned into the pBUTR (piggyBac-based 3′ UnTranslated Region reporter) using gateway cloning as outlined in Chaudhury et al (Chaudhury et al., 2014). Reporters were transfected into ESCs using Fugene 6 (Promega). Cells were treated with Genenticin to enrich for transfected cells. Cells were analyzed on an LSRII (BD). RFP+/GFP+ cells were gated in FlowJo and median RFP/GFP ratios were calculated. RFP/GFP ratios were standardized between days to accounts for differences in laser power.

### Primers

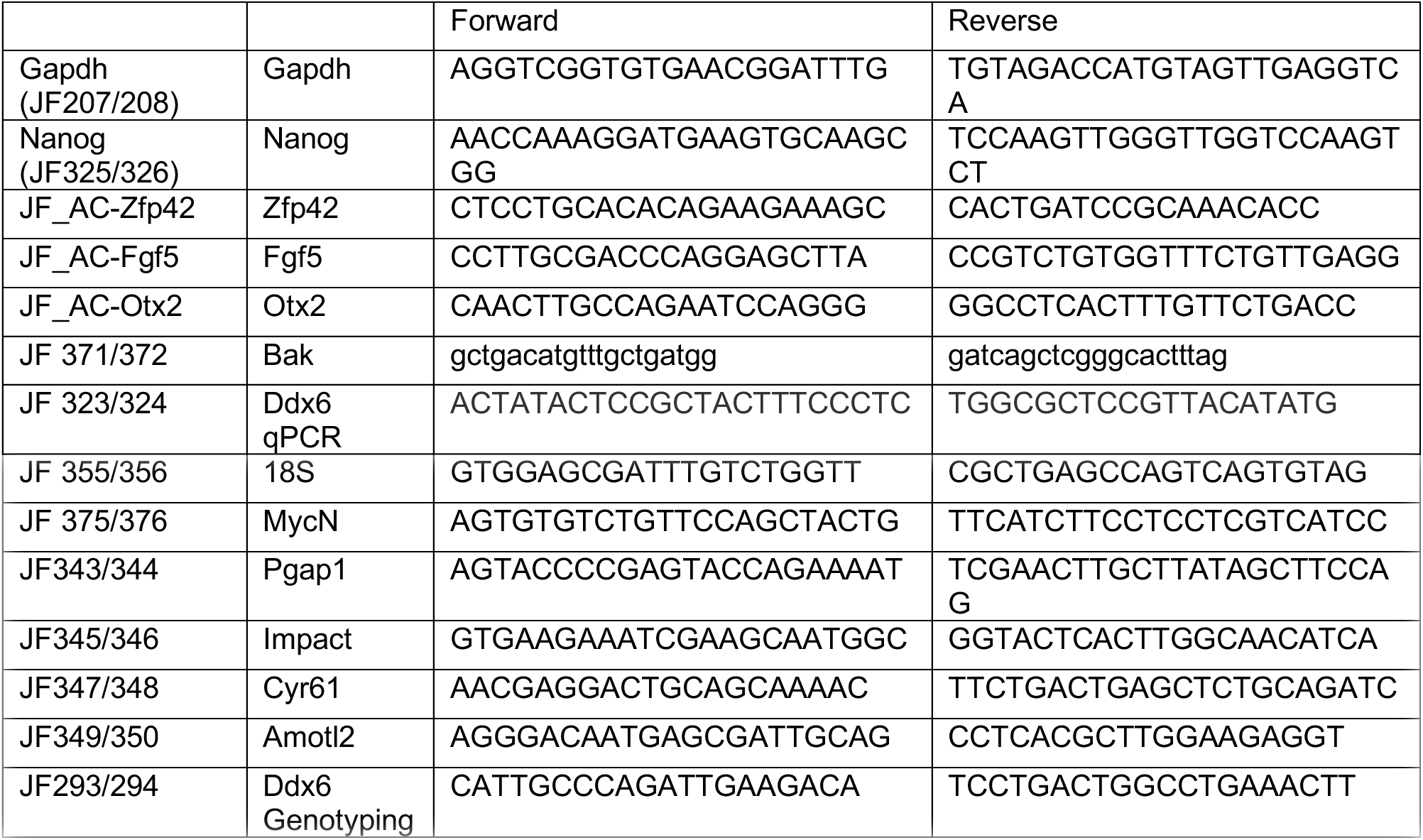

## ACKNOWLEDGEMENTS

We thank the following people for critical reading of the manuscript: Marco Conti, Stephen Floor, Raga Krishnakumar, Brian DeVeale, and Deniz Goekbuget. We also acknowledge Indiana University for access to their Mason cluster of computers, supported by the National Science Foundation (DBI #1458641). This project was funded by the National Institutes of Health (R01 GM101180, R01 GM122439) to R.B., and a Genentech Predoctoral Research Fellowship to J.W.F.

## AUTHOR CONTRIBUTIONS

J.W.F., and R.B. conceived the study, designed the study, and wrote the manuscript. J.W.F. and T.H. performed experiments.

## COMPETING INTERESTS

The authors declare no competing interests.

---

**Figure 1 - figure supplement 1.**
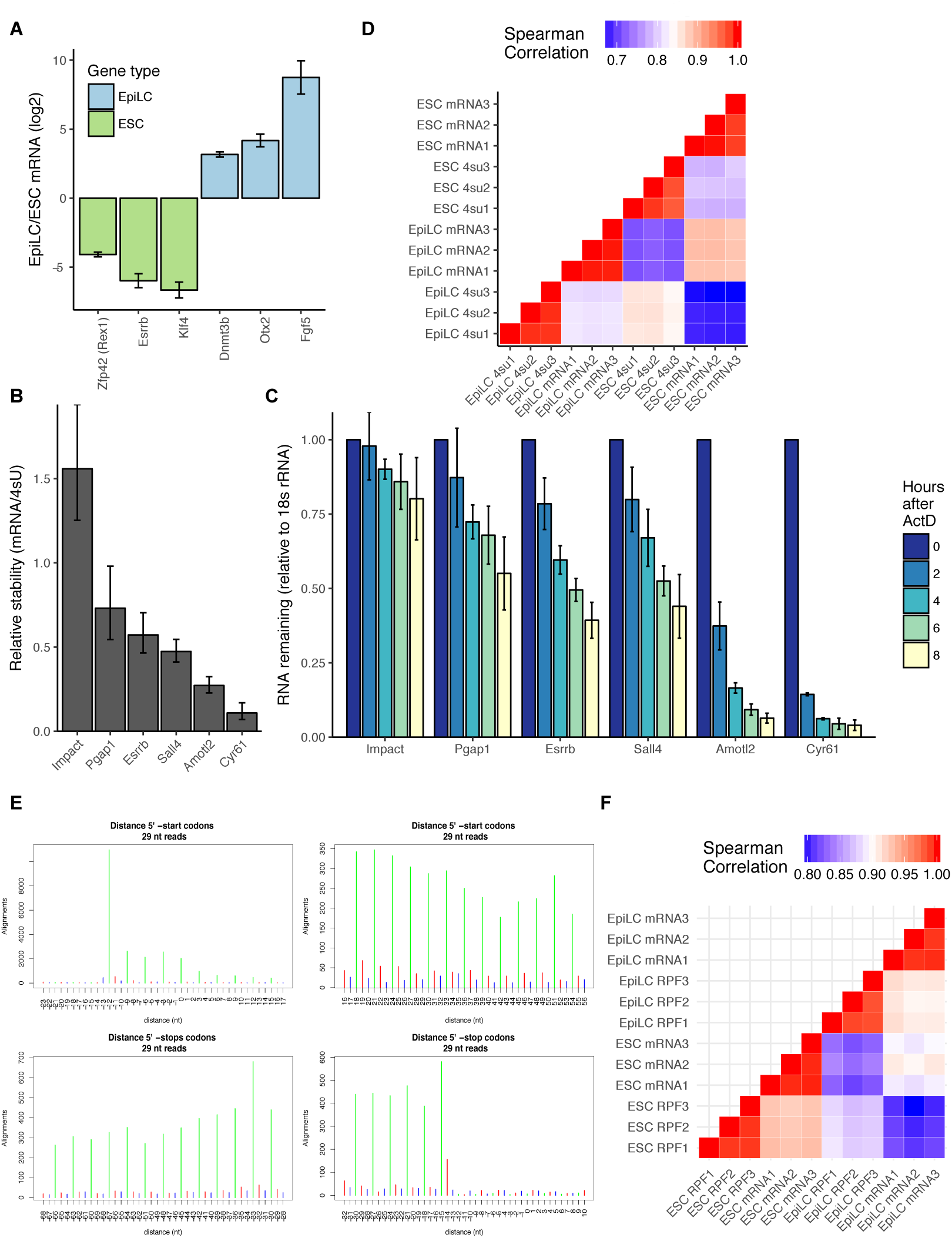
Validation of differentiation, mRNA stability measurements, and ribosome footprinting. A) Change in expression of key naive and primed genes during the ESC to EpiLC transition based on RNA-Seq. Error bars represent 95% confidence interval. B) Relative mRNA stability of candidate genes based on ratio of mRNA/4sU. Error bars represent 95% confidence interval. C) Validation of 4sU-Seq measured mRNA stabilities with RT-qPCR time course after blocking transcription with Actinomycin D. Values are normalized to 18S rRNA and their 0 hr timepoint. n = 3 for wild type and n = 6 (3 replicates of each *Ddx6* KO line), error bars are standard deviation. D) Spearman correlation of log2(counts per million) of ESC and EpiLC RNA-Seq and 4sU-Seq replicates. E) Ribosome profiling shows characteristic phasing for ribosome protected footprints. F) Spearman correlation of log2(counts per million) of ESC and EpiLC RNA-Seq and ribosome protected footprint (RPF) replicates.

**Figure 2 - figure supplement 1.**
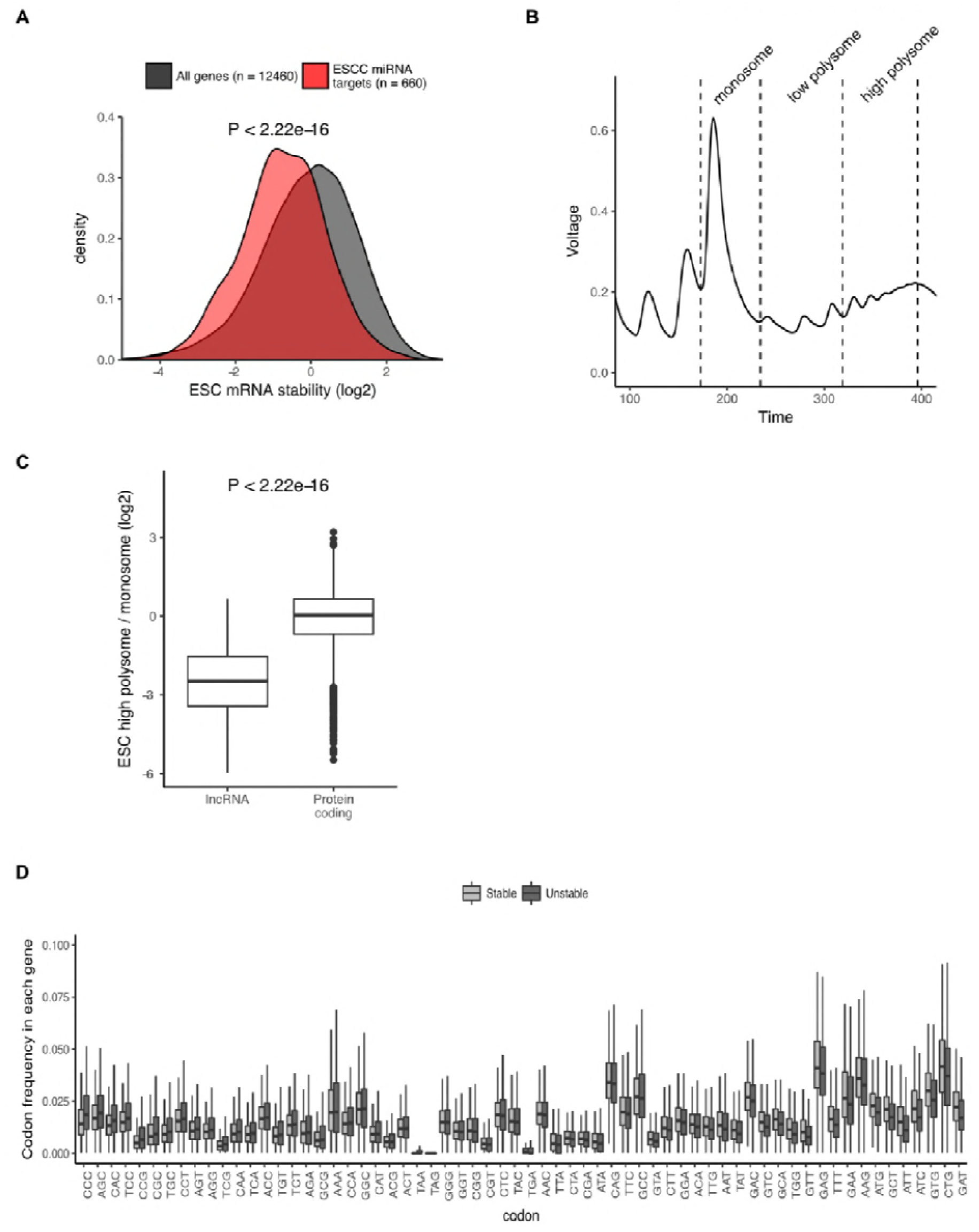
Factors that affect RNA stability in ESCs. A) mRNA stability of ESCC miRNA targets compared to all mRNAs. P value was calculated using the Mann–Whitney test. B) Polysome trace showing monosome, low polysome, and high polysome fractions collected for RNA-Seq. C) Translation level of long non-coding RNAs (lncRNAs) compared to protein coding RNAs. P value calculated with Mann-Whitney test. D) Boxplots showing the codon usage frequency in the top and bottom 20% of genes in terms of stability. Codons are ordered along the X-axis based on the median codon usage in unstable - median codon usage in stable.

**Figure 3 - figure supplement 1.**
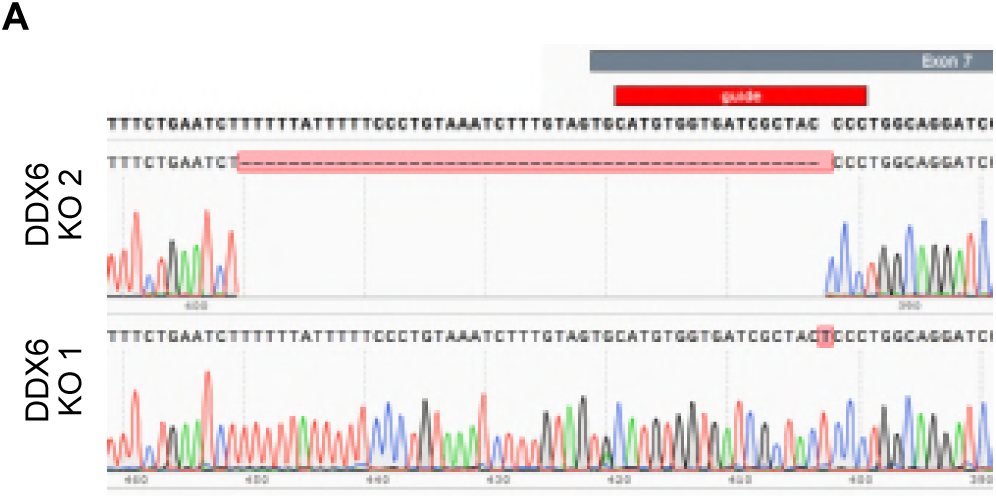
Characterization of *Ddx6* KO ESCs. A) Sanger sequencing showing deletion and insertion at the beginning of exon 7 to generate two *Ddx6* KO clones.

**Figure 4 - figure supplement 1.**
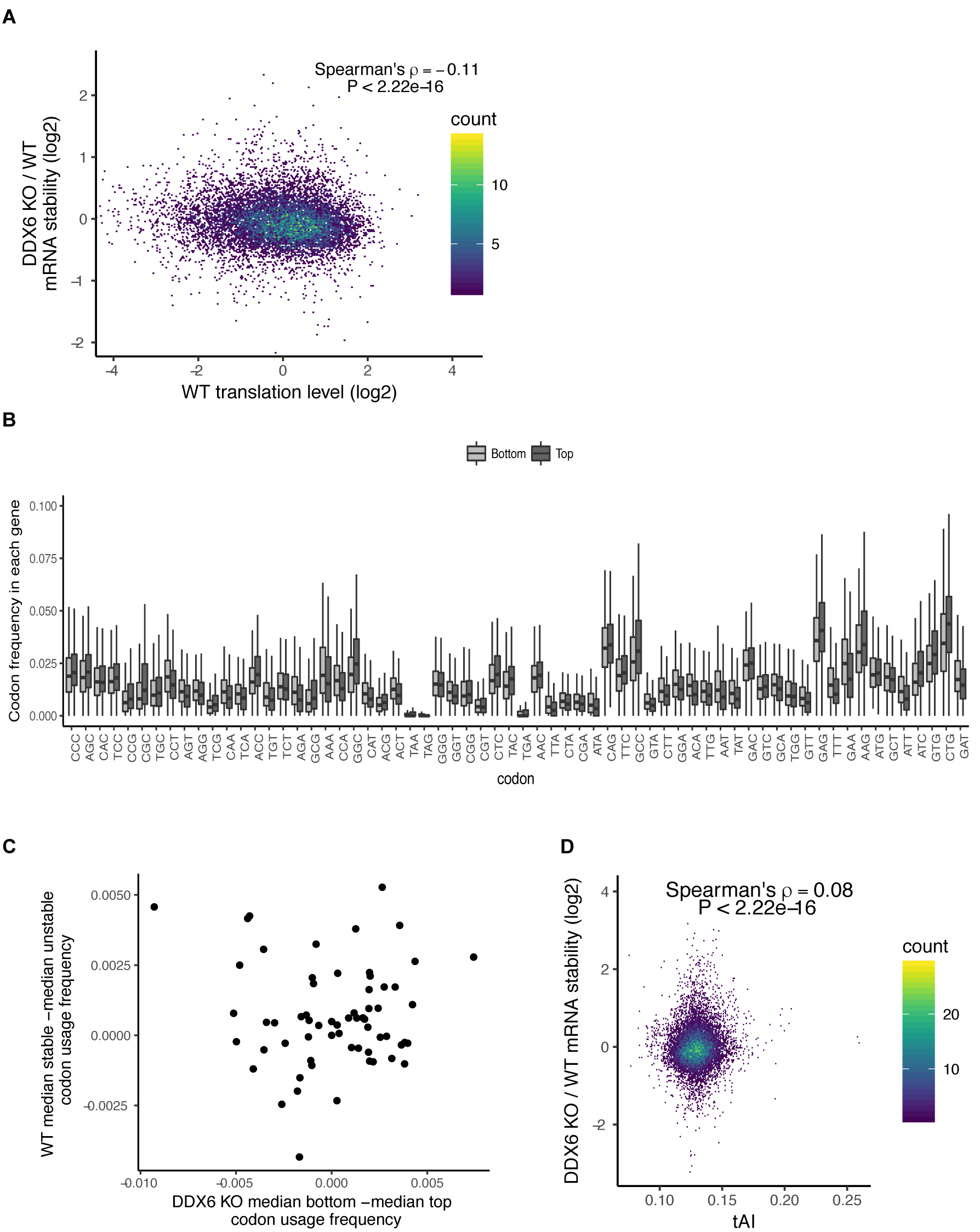
Connection between stability changes and translation. A) mRNA stability changes in *Ddx6* KO cells versus wild type translation level. B) Boxplots showing the codon usage frequency in the top and bottom 20% of mRNA stability changes in *Ddx6* KO for unstable genes as defined in Figure S2D. C) Difference in median codon frequency between stable and unstable transcripts in wild type cells versus difference in median codon frequency between top and bottom *Ddx6* KO mRNA stability changes. D) mRNA stability changes in *Ddx6* KO cells versus species specific tRNA adaptation index (tAI) scores for each gene.

